# Hemin decreases cellular aging and enhances healthspan and lifespan through the AMPK pathway

**DOI:** 10.1101/2023.08.15.553367

**Authors:** Yizhong Zhang, Arshia Naaz, Nashrul Afiq Faidzinn, Sonia Yogasundaram, Trishia Yi Ning Cheng, Jovian Lin Jing, Ingrid Wen-Hui Jeanette Morel Gan, Chen Junqi, Mohammad Alfatah

**Affiliations:** Bioinformatics Institute (BII), Agency for Science, Technology and Research (A*STAR), 30 Biopolis Street, Matrix #07-01, Singapore 138671, Republic of Singapore; Genome Institute of Singapore (GIS), Agency for Science, Technology and Research (A*STAR), 60 Biopolis Street, Genome #02-01, Singapore 138672, Republic of Singapore; Department of Biological Sciences, National University of Singapore (NUS), 16 Science Drive 4, Singapore 117558, Republic of Singapore; Department of Pharmacy, National University of Singapore (NUS), 18 Science Drive 4, Singapore 117543, Republic of Singapore

**Keywords:** Hemin, cellular aging, healthspan, lifespan, AMPK, mitochondria, autophagy, TORC1, yeast, human cells

## Abstract

The quest to understand and manipulate the mechanisms of cellular aging has far-reaching implications for improving human health and longevity. Our comprehensive effort has led to the discovery of the intriguing anti-aging potential of hemin, an FDA-approved drug primarily used for the treatment of acute intermittent porphyria. Leveraging both yeast and human cell models, we investigate the multifaceted effects of hemin on extending cellular lifespan. Intriguingly, the involvement of the AMPK pathway emerges as a pivotal mechanism underlying hemin’s anti-aging effects. The exploration of hemin’s impact on cellular functionality further uncovers its influence on mitochondrial processes. Notably, both mitochondrial-dependent and -independent mechanisms are implicated in hemin’s ability to extend cellular lifespan, with autophagy playing a significant role in the latter. Additionally, a striking synergy between hemin and the TORC1 inhibitor rapamycin is unveiled, underlining the complexity of cellular signaling networks involved in lifespan extension. Translating these findings to human cells, hemin demonstrates an analogous ability to induce mitochondrial biogenesis, reduce proinflammatory cytokine expression, and enhance antioxidant response. The conservation of hemin’s anti-aging effects across species holds promise for therapeutic applications in addressing age-related diseases and promoting healthier aging.

## INTRODUCTION

The world’s aging population is a growing concern, with the proportion of individuals aged 60 and older projected to nearly double from 12% to 22% between 2015 and 2050 ^1^. The global demographic shift towards an aging population has prompted a pressing need for innovative interventions that can enhance healthy longevity and alleviate the escalating burden of chronic diseases such as cancer, cardiovascular diseases, diabetes, neurodegenerative disorders, sarcopenia, and age-related macular degeneration ^2–4^. Cellular aging is a multifaceted process that occurs throughout an individual’s life, encompassing a range of intricate biological and molecular changes ^5–9^. While the resilience of young cells may mask the impairments they have incurred, these underlying cellular dysfunctions can contribute to age-related diseases that manifest later in life ^10^. Furthermore, specific environmental conditions and early-life exposures can expedite the aging process, leading to reduced lifespan and an increased risk of chronic diseases ^11–13^.

Despite being the primary risk factor for numerous human pathologies, cellular aging is often disregarded as a time-dependent physiological phenomenon, or insufficient attention has been given to the role of biological processes in aging-related phenotypes ^14,15^. This limited recognition has resulted in a lack of comprehensive research focused on understanding cellular aging mechanisms. Nevertheless, recent advancements in the field have revealed several promising anti-aging interventions, such as calorie restriction, and specific pharmacological compounds like rapamycin and metformin, which have demonstrated the ability to increase healthspan and lifespan in eukaryotic cells from yeast to humans ^16–24^. Given that cellular aging is a multifaceted process influenced by various molecular pathways ^5–9^, opportunities arise for identifying effective anti-aging compounds capable of slowing down cellular aging without adverse effects, thereby promoting healthier and prolonged lifespans.

In the pursuit of such compounds, yeast, particularly *Saccharomyces cerevisiae*, has emerged as a powerful model organism for unraveling the mysteries of cellular aging and evaluating potential anti-aging interventions ^25–33^. Yeast offers a unique advantage due to its conserved cellular processes and ease of genetic manipulation. Gerontologists have successfully employed two main approaches to study cellular aging in yeast: the replicative lifespan (RLS) assay, which measures the number of cell divisions, and the chronological lifespan (CLS) assay, which quantifies the survival of non-dividing cells. Notably, the CLS assay has gained recognition as a valuable model for studying organismal aging ^26^.

The CLS assay has furnished a comprehensive understanding of the genetic and molecular mechanisms underlying cellular aging. It has facilitated the discovery of the conserved role of several protein complexes and biological processes, including AMP-activated protein kinase (AMPK) and Target of Rapamycin Complex 1 (TORC1), autophagy, and mitochondrial function in cellular aging ^25,26,31,33^. Additionally, the CLS assay has played a pivotal role in investigating the effects of chemical and genetic interventions on cellular aging ^30^, providing insights into potential strategies for promoting healthy aging.

Through this approach, we are evaluating the effects of diverse compounds, including FDA-approved drugs and natural products, on yeast cellular lifespan. Our comprehensive effort has led to the discovery of the intriguing anti-aging potential of hemin, an FDA-approved drug primarily used for the treatment of acute intermittent porphyria ^34^. Our research has unveiled valuable insights into the pivotal role of the AMPK pathway in mediating the anti-aging effects of hemin. Notably, the anti-aging activity of hemin is significantly compromised in mitochondrial and autophagy mutants. Furthermore, our investigations have extended to human cells, demonstrating that hemin effectively extends the healthspan and lifespan of lung fibroblast and kidney cells. This conservation of functional benefits across different species and cell types highlights the promising potential of hemin as a therapeutic agent for anti-aging interventions. Our study has opened a new avenue for further exploration and provided promising leads in the pursuit of enhancing human longevity.

## RESULTS

### Hemin delays cellular aging and extends the lifespan of yeast cells

The utilization of chronological lifespan (CLS) methods allowed us to investigate the impact of hemin on cellular aging. To conduct the CLS experiment, yeast cells were incubated with serial dilutions of hemin ranging from 0.32 µM to 10 µM in the 96-well plate. The CLS was assessed at different time points, providing a comprehensive understanding of the cellular response to hemin treatment. The outgrowth assay revealed striking differences in survival rates between the control and hemin-treated groups. Aged cells without hemin treatment (DMSO control) displayed less than 10% survival rates on day 10 (Figure 1A). In contrast, cells treated with hemin at a concentration of 10 µM exhibited a remarkable increase in survival, reaching approximately 100% on day 10. This enhanced survival was consistently observed at subsequent time points, including day 12, day 16, and day 20 (Figure 1A). These findings strongly support the notion that hemin has the ability to delay cellular aging and extend the lifespan of yeast cells.

**Figure 1.**
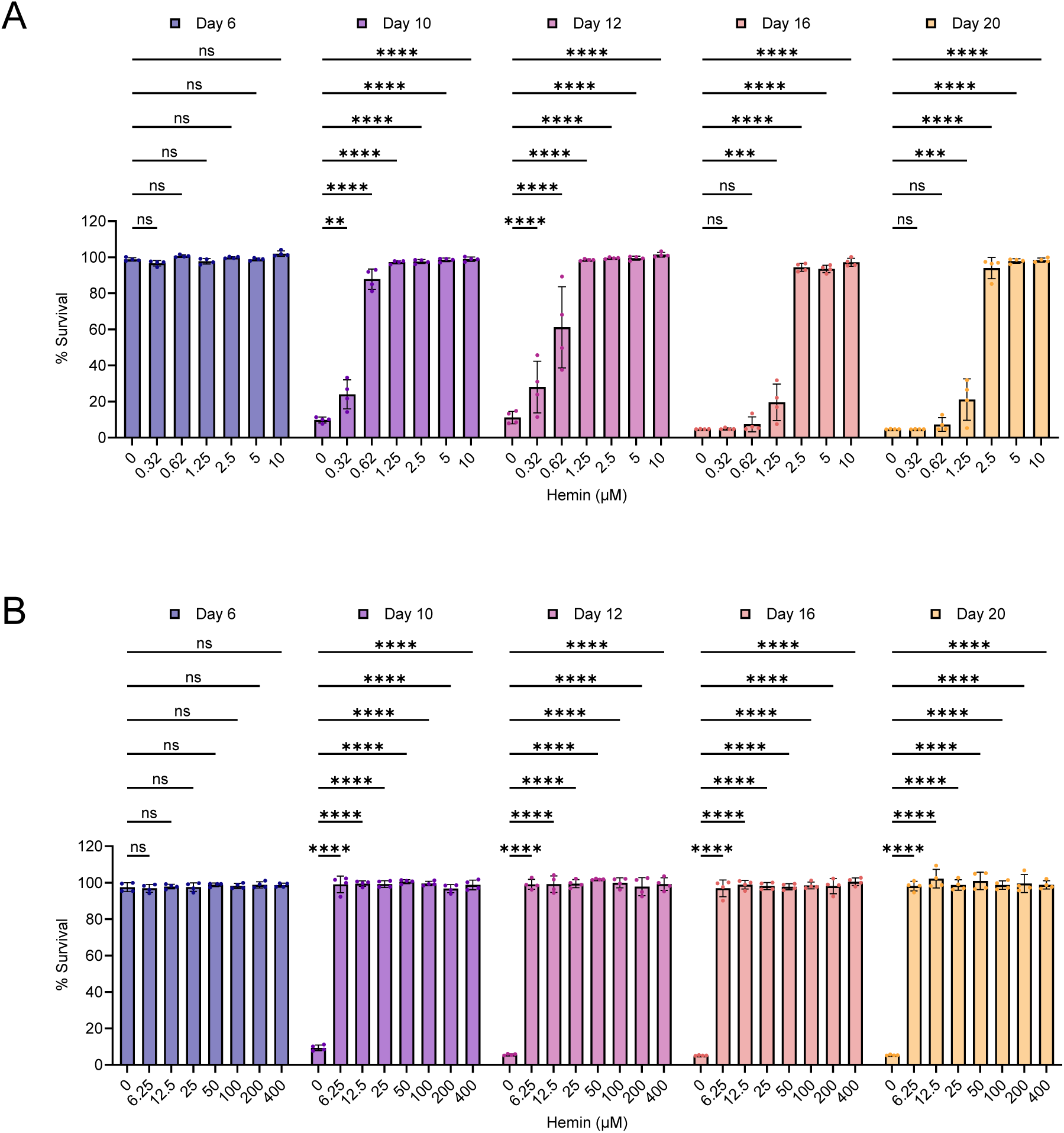
Increased lifespan of yeast cells with hemin treatment. The impact of hemin treatment on the chronological lifespan (CLS) of wild-type yeast cells was evaluated in an SD medium using a 96-well plate. Cells were exposed to varying concentrations of hemin: (A) from 0 µM to 10 µM, and (B) from 0 µM to 400 µM, within the 96-well plate. The survival of aging cells was measured at specified time intervals relative to the outgrowth of day 3. The data are presented as means ± SD (n=4). Statistical significance was determined as follows: **P < 0.01, ***P < 0.001, ****P < 0.0001 and ns: non-significant, based on a two-way ANOVA followed by Dunnett’s multiple comparisons test.

Interestingly, the positive effects of hemin on cellular longevity were not limited to the higher concentration of 10 µM. Even at lower concentrations, hemin demonstrated efficacy in increasing the cellular lifespan. Notably, the lowest concentration tested, 0.62 µM, resulted in approximately 100% survival of aged cells on day 10 (Figure 1A). This intriguing observation suggests that hemin’s anti-aging activity is not solely dependent on high concentrations and that even lower concentrations can significantly impact the cellular lifespan positively.

Considering the biphasic dose-response relationships commonly associated with drug interventions ^35,36^, it was important to determine whether the observed anti-aging effects of hemin followed a similar pattern. We examined the effect of a range of concentrations on cellular lifespan. Serial dilutions of hemin spanning from 6.25 µM to 400 µM were tested for their impact on the CLS of yeast cells. Remarkably, all concentrations of hemin tested resulted in approximately 100% survival of aged cells on day 20 (Figure 1B). These findings indicate that the anti-aging activity of hemin is independent of a hormesis phenotype and can extend cellular lifespan across a wide concentration range.

To further validate the anti-aging properties of hemin, we performed an outgrowth spotting assay on YPD agar medium, which provided a different experimental approach. The cells were incubated with hemin in glass flasks, and the CLS was assessed at different time points. This assay confirmed the consistent and reproducible extension of cellular lifespan associated with hemin treatment (Figure S1A).

We also employed the PICLS method to further validate the anti-aging properties of hemin. In this method, yeast cells were incubated with hemin in a 96-well plate, and the CLS of aging cells was measured using a PI-fluorescence-based assay. The results obtained from the PI-fluorescence assay demonstrated that the presence of hemin significantly increased the survival of aged cells compared to the control-treated cells (Figure S1B).

These additional experimental verifications reinforce the notion that hemin possesses robust anti-aging activity and can effectively prolong the cellular lifespan.

### Anti-aging activity of hemin is dependent on the AMPK pathway

After identifying the role of hemin in cellular longevity, we delved into exploring its anti-aging mechanism. Hemin has demonstrated the ability to activate the AMPK pathway (AMP-activated protein kinase) ^37^. AMPK, a highly conserved master regulator of cellular metabolism ^38,39^, plays a pivotal role in orchestrating diverse biological processes that contribute to cellular homeostasis. Activation of AMPK promotes catabolic processes while inhibiting anabolic pathways, encompassing crucial aspects such as mitochondrial function, autophagy, and the Target of Rapamycin Complex 1 (TORC1) ^39–43^. These interconnected processes governed by AMPK serve as prominent aging hallmarks ^7,8^, influencing the overall cellular aging trajectory ^16,43^.

With cellular aging being closely associated with a progressive decline in AMPK activity, a cascade of events unfolds within cellular machinery, leading to metabolic dysregulation and compromised stress tolerance ^16,43,44^. The cumulative failures and age-dependent inhibitory effect on AMPK activity ultimately culminate in the onset and prevalence of age-related pathologies ^45–48^. Hence, understanding the intricate relationship between hemin and AMPK becomes crucial in elucidating its anti-aging potential.

We aimed to investigate whether the anti-aging activity of hemin is dependent on the AMPK pathway. To achieve this, we turned to the well-established yeast model *Saccharomyces cerevisiae*, in which the Snf1 protein governs AMPK activity ^42,49–51^. First, we sought to evaluate the impact of hemin on AMPK activity in yeast cells. Employing varying concentrations of hemin, we assessed the phosphorylation status of the Snf1 protein. Remarkably, our findings showcased increase in Snf1 protein phosphorylation upon hemin treatment (Figure 2A), corroborating previous studies that have reported the positive influence of hemin on enhancing AMPK activity ^37^.

**Figure 2.**
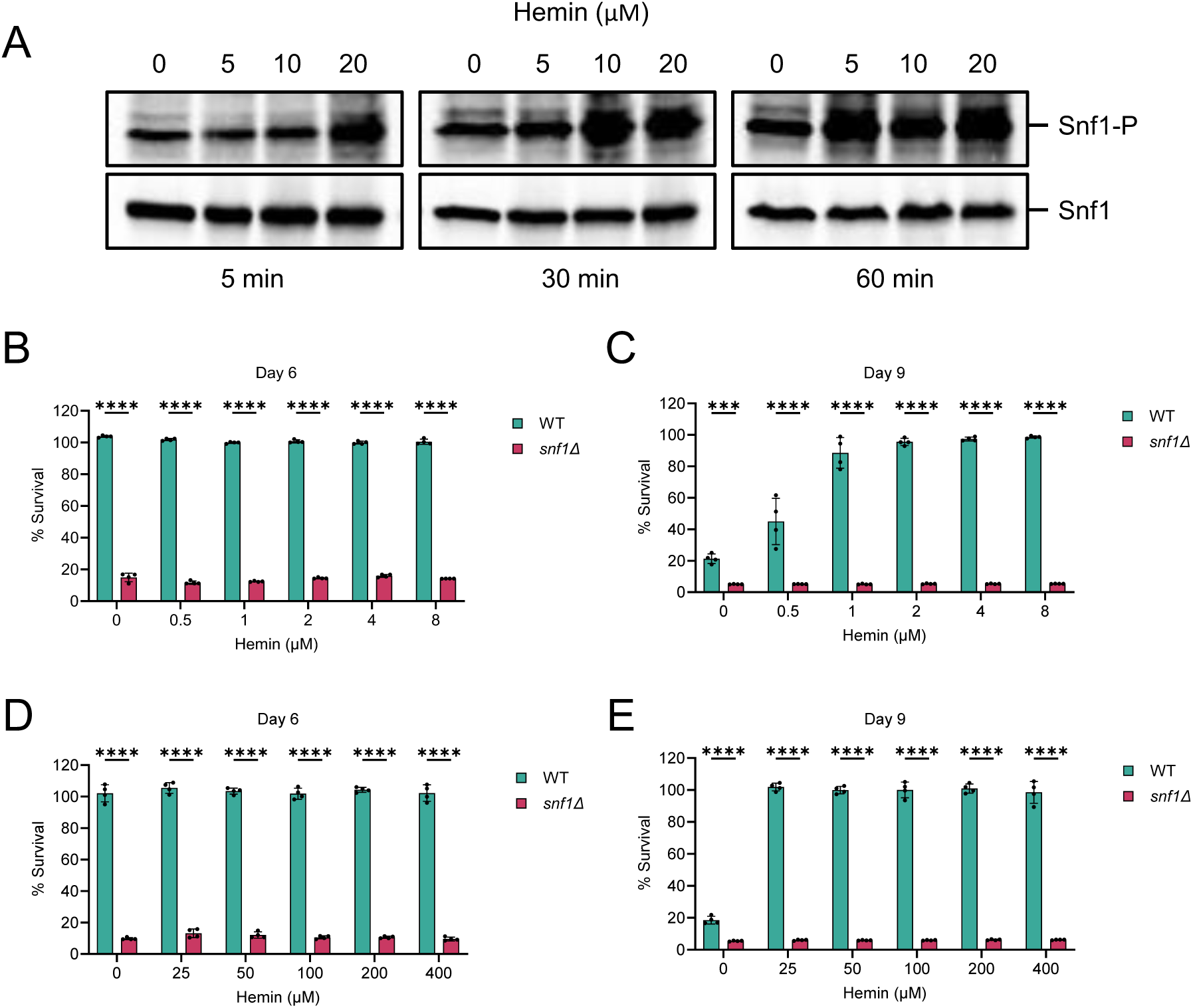
The effectiveness of hemin in increasing cellular lifespan relies on the AMPK pathway. (A) The effect of hemin treatment on AMPK activation was assessed by western blot analysis of phosphorylated Snf1 using anti-phospho-AMPKα antibody. Lower blots were probed with anti-MYC-Tag antibody to detect total Snf1 protein. Cells were obtained from a wild-type strain (Snf1-9xMyc-tag) with exponential cultures in SD medium, treated with varying concentrations of hemin for the indicated durations. (B-E) The impact of hemin treatment on the chronological lifespan (CLS) of wild-type and AMPK-deficient (*snf1Δ* deletion) yeast cells was evaluated in SD medium using a 96-well plate. Cells were exposed to lower (B and C) and higher (D and E) concentrations of hemin within the 96-well plate. Survival of aging cells was measured at specified time intervals relative to the outgrowth of day 3. The data are presented as means ± SD (n=4). Statistical significance was determined as follows: ***P < 0.001, and ****P < 0.0001, based on a two-way ANOVA followed by Šídák’s multiple comparisons test.

To further elucidate the connection between hemin, AMPK, and cellular aging, we examined the effect of hemin on the lifespan of *SNF1*-deleted cells. Notably, the deletion of the *SNF1* gene (*snf1Δ*) is known to decrease the lifespan of wild-type cells (Figure S2) ^52,53^, thereby implicating its functional role in cellular aging. Building upon this observation, we conducted lifespan experiments on *snf1Δ* mutants treated with varying concentrations of hemin (ranging from 0.5 µM to 8 µM). Intriguingly, our results unveiled a significant disparity: hemin treatment failed to extend the chronological lifespan (CLS) of *snf1Δ* cells (Figure 2B and 2C), suggesting that the anti-aging activity of hemin is indeed dependent on the AMPK pathway.

Given that lower concentrations were unable to increase the cellular lifespan of the *snf1Δ* mutant, we hypothesized that higher concentrations might be necessary to achieve this effect. Therefore, we investigated the CLS of *snf1Δ* cells that were incubated with varying concentrations of hemin, ranging from 25 µM to 400 µM. Surprisingly, even at these higher concentrations, hemin was ineffective in enhancing the cellular lifespan of *snf1Δ* mutant (Figure 2D and 2E). These findings align consistently with the CLS phenotype observed in *snf1Δ* cells exposed to lower concentrations of hemin (Figure 2B and 2C). Altogether, our comprehensive results emphasize the essential role of AMPK activity for hemin to significantly enhance cellular lifespan, thereby providing novel insights into the intricate interplay between hemin, AMPK, and the aging process.

### Hemin increases cellular lifespan via mitochondrial function

One of the downstream effectors of the AMPK pathway is mitochondrial function ^38,39^. AMPK is activated in response to a decrease in cellular energy, triggered by changes in the ATP-to-ADP or ATP-to-AMP ratio. The allosteric activation of AMPK stimulates its kinase activity, leading to metabolic reprogramming towards increased catabolic pathways ^16,38–40,54^. Interestingly, mitochondrial functions also decline with cellular aging and are implicated in age-related diseases ^16,44,55–57^.

Given the well-established relationship between hemin’s anti-aging effects and AMPK, we conducted a study to explore its impact on mitochondrial functions ^58^. To initiate our investigation, we evaluated the effect of hemin on cellular energy levels. Notably, our results demonstrated that treatment with hemin leads to a notable increase in cellular ATP levels (Figures 3A; S3A). Furthermore, we focused on the Electron Transport Chain (ETC) within the mitochondria, which plays a pivotal role in generating ATP through oxidative phosphorylation (OXPHOS) processes ^58^. Intriguingly, the expression of specific ETC genes exhibited an upregulation following hemin treatment (Figure 3B; S3B), indicating a potential enhancement in ATP production. A key player in the regulation of mitochondrial biogenesis is the Hap4 transcription factor, responsible for orchestrating the expression of specific ETC genes ^59^. It is worth noting that the expression of *HAP4* was also notably elevated in cells treated with hemin (Figures 3B; S3B). Collectively, these findings suggest that hemin treatment instigates a remodeling of cellular metabolism. This restructuring is achieved through the promotion of mitochondrial gene expression and the augmentation of energy synthesis pathways.

**Figure 3.**
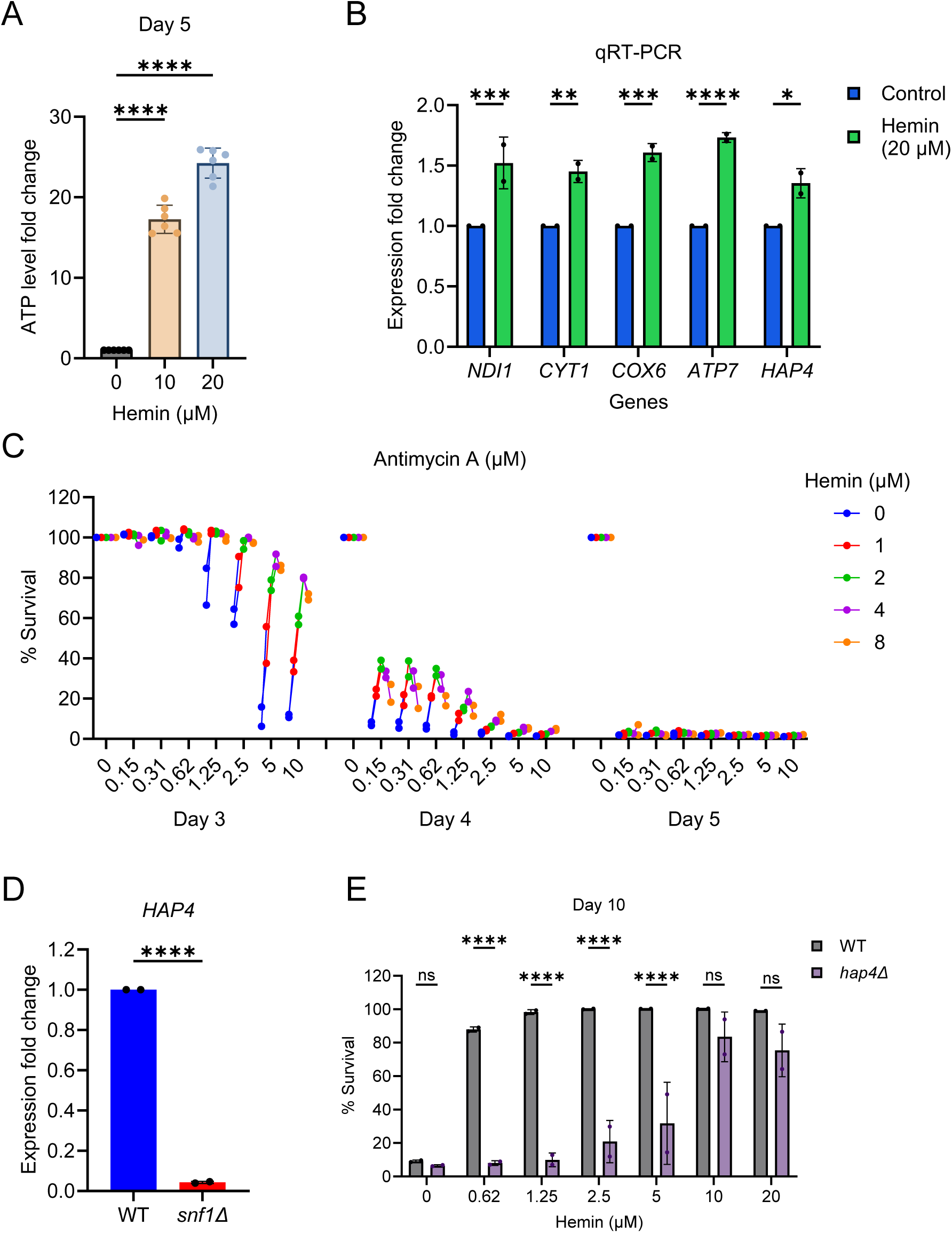
Hemin extends cellular lifespan via mitochondrial function-dependent and independent mechanisms. (A) ATP analysis of wild-type yeast cells incubated with different concentrations of hemin for 5 days. The data are presented as means ± SD (n=6). ****P < 0.0001, based on an ordinary one-way ANOVA followed by Dunnett’s multiple comparisons test.. (B) Expression analysis of mitochondrial ETC genes by qRT-PCR in yeast cells incubated with 20 µM hemin for 6 hours, starting at ∼0.2 OD600 nm. The data are presented as means ± SD (n=2). Statistical significance was determined as follows: *P < 0.05, **P < 0.01, ***P < 0.001, and ****P < 0.0001 based on a two-way ANOVA followed by Šídák’s multiple comparisons test. (C) Chronological lifespan (CLS) of wild-type yeast strain with varying concentrations of antimycin A (0 µM - 10 µM) and hemin (0 µM - 8 µM) was assessed in SD medium in a 96-well plate. Survival of aging cells was measured relative to the control outgrowth for the indicated day of analysis. The data are represented from two experiments. (D) Expression analysis of the *HAP4* gene by qRT-PCR in exponentially growing wild-type and AMPK-deficient (*snf1Δ* deletion) yeast cells. The data are presented as means ± SD (n=2). Statistical significance was determined as follows: ****P < 0.0001 based on a two-sided Student’s t-test. (E) Chronological lifespan (CLS) of wild-type and *hap4Δ* deletion yeast cells with indicated concentrations of hemin was assessed in SD medium in a 96-well plate. Survival of aging cells was measured at specified time intervals relative to the outgrowth of day 3. The data are presented as means ± SD (n=2). Statistical significance was determined as follows: ****P < 0.0001 and ns: non-significant, based on a two-way ANOVA followed by Šídák’s multiple comparisons test.

To investigate the coupling of hemin’s effects with cellular aging, we examined the chronological lifespan (CLS) of dysfunctional mitochondrial cells. We used Antimycin A (AMA), an inhibitor of the mitochondrial ETC pathway ^60^. Upon treating wild-type cells with different concentrations of AMA, we evaluated the CLS. Concurrently, we co-administered hemin with AMA during the chronological aging process and monitored cell survival. The results demonstrated that AMA treatment reduced the lifespan of wild-type cells (Figure 3C), confirming the regulatory role of mitochondrial functions in cellular aging. On day 3, hemin supplementation significantly rescued shortened lifespan of AMA-treated cells, partially on day 4 (Figure 3C). However, noteworthy, on day 5, hemin failed to increase the lifespan of AMA-treated cells (Figures 3C). These observations suggest that hemin affects cellular aging through both mitochondrial function-dependent and -independent mechanisms.

Hap4 activity is associated with extending cellular lifespan by enhancing mitochondrial function ^59^. To understand the interplay between the AMPK pathway and Hap4 activity, we examined whether the AMPK pathway influences Hap4 activity. Our findings revealed that *HAP4* gene expression was downregulated in *snf1Δ* cells (Figure 3D), indicating a link between the AMPK pathway and Hap4 activity. Additionally, we investigated the effect of Hap4 activity on hemin’s impact on cellular lifespan. We found that the anti-aging effect of hemin was significantly compromised in *hap4Δ* mutant cells (Figure 3E). This observation aligns with our previous findings, suggesting that hemin’s anti-aging effect is dependent on both mitochondrial function coupled with AMPK activity and independent mechanisms.

### Hemin increases cellular lifespan via autophagy-dependent and TORC1-independent mechanisms

To explore the mitochondrial-independent mechanism responsible for hemin’s anti-aging activity, we embarked on an investigation of alternative effectors within the AMP-activated protein kinase (AMPK) pathway. While we had previously highlighted AMPK’s role in enhancing mitochondrial function, it is also known to positively regulate the autophagy process ^43,61,62^, a fundamental cellular mechanism for recycling and degrading cellular components. One intriguing form of autophagy is mitophagy, a selective process that specifically targets and degrades damaged or dysfunctional mitochondria. This remarkable process plays a crucial role in preventing accelerated aging and promoting extended lifespan in various organisms ^54,55,57^.

Driven by the need to understand the connection between hemin and the autophagy process, we delved deeper into our investigation. To shed light on hemin’s anti-aging activity and its dependence on autophagy, we conducted lifespan experiments using specific autophagy mutants (*atg1Δ* and *atg7Δ*) ^63^ and a mitophagy-specific mutant (*atg32Δ*) ^64^, in addition to wild type and *snf1Δ* mutant cells.

The results of our study corroborated earlier findings, demonstrating consistent phenotypic patterns of hemin’s impact on both wild type and *snf1Δ* mutant cells (Figures 4A and 4B). However, we observed that the effect of hemin on the lifespan of autophagy mutants was notably reduced compared to its impact on wild type cells (Figures 4C - 4E). This compelling evidence strongly suggests that hemin’s ability to increase cellular lifespan is intrinsically linked to an autophagy-dependent mechanism.

**Figure 4.**
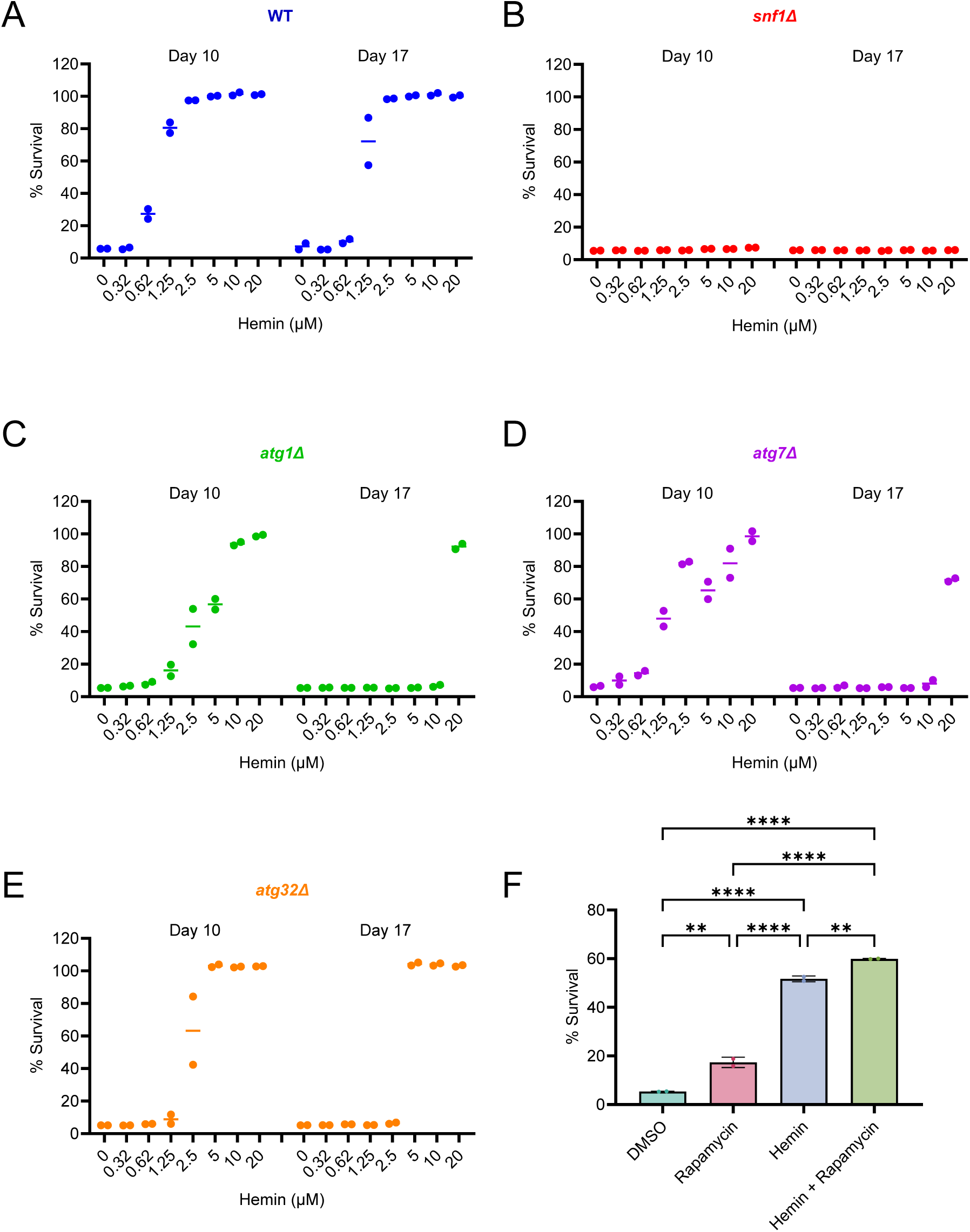
Hemin extends cellular lifespan through autophagy-dependent and TORC1 -independent mechanisms. (A-E) Evaluation of the Impact of hemin treatment on chronological lifespan (CLS) in wild-type, AMPK-deficient, and autophagy-deficient yeast cells using a 96-well plate assay in SD medium. Various concentrations of hemin were incubated with (A) wild-type, (B) *snf1Δ*, (C) *atg1Δ*, (D) *atg7Δ*, and (E) *atg32Δ* strains within a 96-well plate. Survival of aging cells was measured at specified time intervals relative to the outgrowth of day 3. Data represent two experiments. (F) CLS analysis of wild-type cells treated with DMSO control, rapamycin (2 nM), hemin (2 µM), and a combination of rapamycin (2 nM) and hemin (2 µM) in SD medium using flask cultures. Outgrowth of aging cells was performed on YPD medium. Survival of aging cells was measured on day 6 relative to the outgrowth of day 3. Data are presented as means ± SD (n=2). Statistical significance was determined as follows: **P < 0.01, and ****P < 0.0001, based on an ordinary one-way ANOVA followed followed by Tukey’s multiple comparisons test.

With the knowledge that the target of rapamycin complex 1 (TORC1) negatively regulates the autophagy process ^40,61–63,65^, we sought to explore the role of TORC1 in mediating the anti-aging effects of hemin. Rapamycin, a well-known drug, is recognized for its ability to inhibit TORC1 activity, resulting in an extension of cellular lifespan across different species, from yeast to human cells ^66–70^. We designed a series of experiments to scrutinize the cellular lifespan of yeast cells treated with hemin alone, rapamycin alone, and a combination of both compounds.

As expected, both hemin and rapamycin treatments independently contributed to an increased cellular lifespan (Figure 4F). However, our most intriguing observation was made when combining hemin and rapamycin. The results revealed a synergistic effect, as the combination treatment further extended the lifespan of cells compared to individual treatments with either hemin or rapamycin (Figure 4F). This remarkable finding strongly suggests that hemin’s anti-aging activity is not only independent of TORC1 but also has the potential to interact beneficially with the TORC1 pathway, further enhancing cellular lifespan.

To solidify our conclusions, we took an additional step to directly assess hemin’s impact on TORC1 activity by monitoring the phosphorylation status of the substrate Sch9 through western blot analysis ^71^. As expected, rapamycin, a potent TORC1 inhibitor, successfully suppressed TORC1 activity, as evidenced by reduced Sch9 phosphorylation (Figure 4SA). Intriguingly, in hemin-treated cells, TORC1 activity remained comparable to that of control untreated cells (Figure 4SB). These critical findings provide compelling evidence that hemin’s anti-aging activity operates independently of the TORC1 signaling pathway, further corroborating our earlier conclusions.

Taken together, our comprehensive investigation into the mitochondrial-independent mechanism underlying hemin’s anti-aging activity has shed new light on the intricate interplay between AMPK, autophagy, and TORC1 pathways. Hemin’s ability to extend cellular lifespan through autophagy-dependent mechanisms highlights its potential aging-related processes.

### Hemin increases the healthspan and lifespan of human cells

The preceding discoveries within the context of the yeast aging model organism have provided compelling evidence that underscores the capacity of hemin to effectively delay the aging process and promote increased longevity at the cellular level. To further advance our understanding, we embarked on investigating whether the favorable attributes of hemin’s impact are preserved within the intricate cellular machinery of human cells.

Our investigation has uncovered that the anti-aging activity of hemin within yeast axes upon the AMP-activated protein kinase (AMPK) pathway. This pathway exerts its influence by orchestrating the modulation of the Hap4 transcription factor, which subsequently augments mitochondrial functionality. Drawing parallels to human cellular dynamics, we assessed the influence of hemin on mitochondrial biogenesis. In the human cells, the peroxisome-proliferator-activated receptor gamma coactivator 1α (PGC-1α) stands as the master regulator governing mitochondrial biogenesis ^72^. Notably, the AMPK pathway orchestrates the expression of PGC-1α ^73,74^. By subjecting hemin-treated human fibroblast cells (IMR90) ^75^, we have ascertained a marked elevation in PGC-1α expression (Figures 5A; S5A).

**Figure 5.**
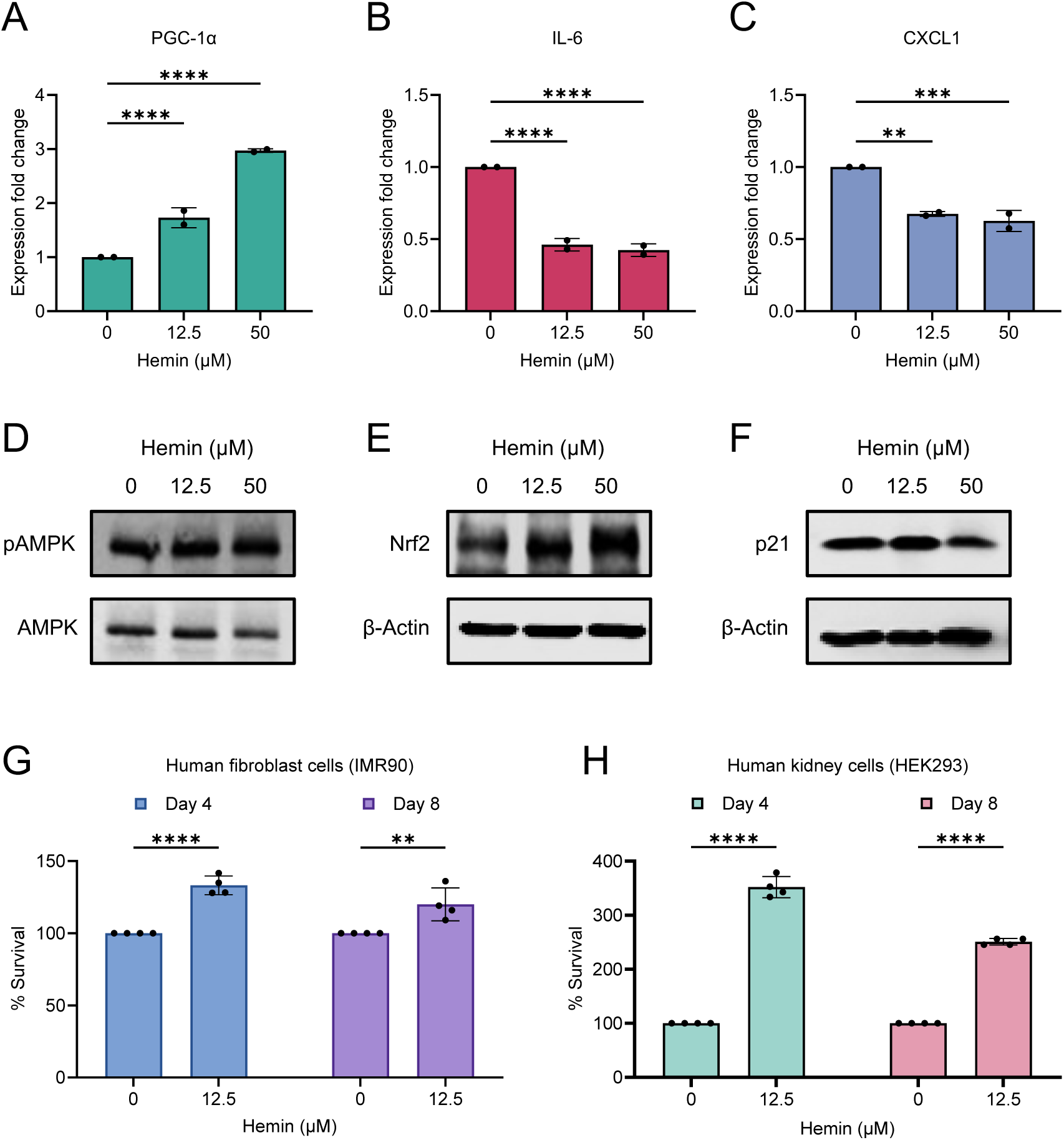
Hemin enhnaces healthspan and lifespan of human cells. (A-C) Expression analysis of gene (A) PGC-1α, (B) IL-6 and (C) CXCL1 by qRT-PCR in human lung fibroblast cells (IMR90) treated with indicated concentrations hemin for 6 hours. The data are presented as means ± SD (n=2). Statistical significance was determined as follows: **P < 0.01, ***P < 0.001, and ****P < 0.0001 based on a two-way ANOVA followed by Dunnett’s multiple comparisons test. (D-F) The effect of hemin treatment on (D) activity of AMPK activity, and expression of (E) Nrf2 and (F) p21 was assessed by western blot analysis. Cells were obtained from IMR90 cells treated with indicated concentrations of hemin for the 4 hours. (G and H) Chronological lifespan (CLS) determination of (G) human lung fibroblast cells (IMR90) and (H) human kidney cells (HEK293) cells with indicated concentrations of hemin was assessed in D10 medium in a 96-well plate. Survival of aging cells was measured relative to the control outgrowth for the indicated day of analysis. Data are presented as means ± SD (n=4). Statistical significance was determined as follows: **P < 0.01, and ****P < 0.0001, based on a two-way ANOVA followed by Šídák’s multiple comparisons test.

A pivotal marker of healthspan, PGC-1α, plays a multifaceted role encompassing anti-inflammatory mechanisms by inhibiting the expression of proinflammatory cytokines and inducing the cellular defenses against oxidative stress ^76,77^. Within this paradigm, interleukin 6 (IL-6) emerges as a prominent proinflammatory factor implicated in propagating inflammatory cascades that ultimately undermine cellular healthspan ^76–78^. Given hemin’s capacity to augment PGC-1α expression, our inquiry extended to discerning its impact on IL-6. Encouragingly, our findings underscored that hemin administration precipitates a reduction in IL-6 expression (Figures 5B; S5B). Concomitantly, hemin-treated cells exhibited a discernible reduction in the expression of CXCL1 (Figures 5C; S5C), a pivotal chemokine that exercises regulatory control over inflammatory responses ^79^. The cumulative outcome of these observations serves to underline hemin’s efficacy in enhancing biological markers intricately linked to healthspan.

In a coherent reflection of these revelations, we probed the impact of hemin treatment on the AMPK activation (Figure 5D) and the expression of the antioxidant transcription factor Nrf2 (nuclear factor-like 2) (Figure 5E) ^48,80,81^. Furthermore, we investigated Nrf2-dependent p21 senescence marker ^80^, which exhibited a reduction within hemin-treated cells (Figure 5F). To conclusively establish the healthspan-boosting attributes of hemin, our focus shifted to assessing cellular survival under conditions simulating chronological aging. Notably, our investigations affirmed a significant augmentation in the lifespan of fibroblast cells upon hemin treatment (Figure 5G).

To broaden the scope, we probed the impact of hemin on the lifespan of an alternative cell type, namely kidney cells (HEK293). Significantly, the extension of lifespan observed in kidney cells upon hemin treatment mirrored the outcomes witnessed in fibroblast cells (Figure 5H). Collectively, these findings coalesce to accentuate that hemin’s anti-aging potential remains robustly conserved across diverse species and distinct human cell lineages.

## DISCUSSION

In this cellular aging context, the innovative research presented in this study offers a promising solution for healthspan. Through a detailed investigation into the effects of hemin on cellular aging and longevity, we have unveiled a potential geroprotective intervention that holds significant promise for the aging population. Our findings reveal that hemin, an FDA-approved drug ^34^, possesses robust anti-aging properties, effectively delaying cellular aging and extending cellular lifespan.

Our investigation explored into the mechanisms underpinning hemin’s anti-aging effects, revealing a detailed and multifaceted framework orchestrated through pivotal intracellular pathways. At the core of this framework lies the AMPK pathway, a master regulator of cellular energetics ^16,38,39,54,82^. The activation of AMPK emerged as a key nexus in mediating hemin’s anti-aging impact. A critical aspect of our exploration entailed a detailed examination of *SNF1*-deleted (*snf1Δ*) yeast cells. The absence of *SNF1* resulted in a decreased lifespan of wild-type cells, unequivocally implicating its indispensable role in the continuum of cellular aging. Intriguingly, the introduction of hemin yielded a complete abolishment in extending the lifespan of *snf1Δ* cells, firmly establishing the AMPK pathway as an essential conduit for hemin’s anti-aging effects.

AMPK is intricately associated with initiating a cascade of events that culminate in the augmentation of mitochondrial bioenergetics, the orchestration of autophagy, and a profound interplay with the TORC1 pathway ^39–43^. To further unravel the role of AMPK effectors in hemin’s anti-aging activity, we initially investigated its impact on mitochondrial functions ^58^. Hemin intervention catalyzed a notable escalation in cellular ATP levels, underscoring a substantial amplification of energy generation. This heightened state of cellular energy resonated within the expression patterns of ETC genes, implying a potential enhancement in ATP production through mitochondrial OXPHOS systems ^58^. Notably, the prominent Hap4 transcription factor ^59^, a key player in mitochondrial biogenesis, emerged as a central figure in this dynamic interplay. Hemin treatment induced a marked increase in *HAP4* gene expression, leading to an intricate recalibration of cellular energetics. In this reconfigured landscape, hemin facilitated an augmented flux through energy synthesis pathways, accompanied by a commendable enhancement in mitochondrial competence. However, it is noteworthy that the capacity of hemin to extend lifespan is significantly compromised in cells harboring chemically induced dysfunctional mitochondria or in *hap4Δ* mutant cells. These observations collectively suggest that hemin exerts its effect on cellular aging through a complex interaction of mechanisms that involve both mitochondrial function-dependent and -independent pathways.

Expanding our investigation, we turned our attention to another AMPK effector, autophagy, an evolutionarily conserved process of intracellular catabolism and turnover ^43,61,62^. The intricate interaction between hemin and autophagy came to light through lifespan experiments involving autophagy-deficient mutants. These observations underscored the indispensable role of autophagy-mediated mechanisms in mediating hemin’s anti-aging effects. Intriguingly, the TORC1 pathway, a pivotal regulator of autophagy modulation ^40,61–63,65^, followed a distinct trajectory from hemin’s orchestrated effects. The notable synergy in the extension of cellular lifespan observed upon concurrent administration of hemin and rapamycin provided intriguing insights into hemin’s potential interactions with the TORC1 pathway, revealing an intricate interplay that warrants further exploration.

Transcending the confines of yeast cells, our exploration extended into human fibroblast and kidney cells, unveiling a remarkable conservation of hemin’s anti-aging efficacy across disparate cellular lineages. Hemin administration prompted an elevation in PGC-1α expression, a pivotal regulator of mitochondrial biogenesis within human cells ^72^. This enhanced expression synchronized with a discernible attenuation in the manifestation of proinflammatory mediators, thereby highlighting hemin’s beneficial impact on healthspan features.

Notably, hemin’s effects extended to cytokine dynamics within human cells. Analysis of interleukin 6 (IL-6), a prominent proinflammatory factor ^76–78^, revealed a notable reduction in its expression following hemin administration, underscoring hemin’s potential in attenuating age-associated inflammaging and modulating cellular resilience. Concomitantly, the expression of CXCL1, a pivotal chemokine that effects regulatory control over inflammatory responses ^79^, was also notably diminished upon hemin treatment. Furthermore, our investigation uncovered hemin’s interaction with the antioxidant transcription factor Nrf2, a sentinel guardian against oxidative stress ^48,80,81^. Expression of Nrf2 was evidently enhanced in the milieu of hemin treatment, indicative of a coordinated response aimed at strengthening cellular resilience against oxidative insults.

Collectively, our study presents a thorough demonstration of hemin’s anti-aging effects, unveiling the intricate interplay of cellular pathways and orchestrations that culminate in its effects. We discovered the novel role of hemin in anti-aging activity that decreases the cellular aging and extends healthspan via AMPK pathway axis-mitochondria and autophagy. These findings underscore hemin’s potential as a geroprotective adjunct, offering a nuanced and multi-pronged strategy to elevate healthspan and extend lifespan. By targeting cardinal mechanisms pivotal to cellular aging, hemin emerges as a captivating candidate for further research and practical applications. This shines a hopeful light on the potential for people to enjoy longer lives with better quality as they grow older.

## METHODS

### Yeast strains and growth conditions

The *S. cerevisiae* prototrophic CEN.PK113-7D strains were used in this study ^83^. Gene deletion and protein tagging was generated using standard PCR-based method ^84^. Yeast strains were revived from frozen glycerol stock on YPD agar (1% Bacto yeast extract, 2% Bacto peptone, 2% glucose and 2.5% Bacto agar) medium for 2-3 days at 30°C.

### Human cell lines and growth conditions

Human lung fibroblast cell line (IMR-90) (Coriell Institute, Camden, New Jersey, United States of America) and human embryonic kidney cell line (HEK293) (ATCC, Manassas, Virginia, United States of America) were cultured in the standard D10 medium, consisting of high-glucose DMEM (HyClone #SH30022.01) supplemented with 10% FBS (Gibco™ #10270106) and 1% Penicillin Streptomycin Solution (Gibco™ #15140122). All cells were cultured in a humidified incubator with 5% CO2 at 37°C.

### Chemical treatment to cell culture

The stock solution of hemin, antimycin A, and rapamycin was prepared using dimethyl sulfoxide (DMSO) as the solvent. The ultimate concentration of DMSO remained below 1% for yeast experiments and below 0.01% for experiments involving human cell lines.

### Chronological lifespan analysis of yeast cells

Chronological Lifespan (CLS) was assessed by determining the survival of aging cells, as previously described ^30^. Yeast cultures were grown in synthetic defined (SD) medium (6.7 g/L yeast nitrogen base with ammonium sulfate without amino acids and 2% glucose) overnight at 30°C with 220 rpm shaking. The cultures were then diluted to a starting optical density at 600 nm (OD600) of approximately 0.2 in fresh SD medium to initiate the CLS experiment. The CLS experiment employed three methods: (i) outgrowth in YPD medium, (ii) outgrowth by spotting on YPD agar, and (iii) PICLS. In brief, cell cultures were grown and aged in 96-well plates or glass flasks in SD medium at 30°C. At various chronological time points, the survival of aging cells was determined by measuring the outgrowth (OD600nm) in YPD medium incubated for 24 hours at 30°C, using a microplate reader. For cells grown in flasks at different age time points, cultures were washed and normalized to an OD600nm of 1.0. Yeast cells were serially 10-fold diluted and 3 μL of the dilution was spotted onto YPD agar plates, followed by incubation for 48 hours at 30°C. The outgrowth of aged cells on the YPD agar plate was documented using the GelDoc imaging system. In the PICLS method, the survival of cells at various chronological age time points was determined using a PI fluorescence-based assay. PI fluorescence reading (excitation at 535nm, emission at 617nm) and OD600nm were measured using a microplate reader. The fluorescence intensity of each sample was normalized to OD600nm.

### Chronological lifespan analysis of human cells

The chronological lifespan (CLS) of human cells was evaluated by determining the survival of aging cells using the outgrowth method. Cells (8 x 104) were initially seeded into a 96-well culture plate with varying concentrations of hemin and incubated for aging at 37°C with 5% CO2. At different chronological time points, aliquots of 5% or 10% of the cells were transferred to 96-well experimental plates containing fresh D10 medium for the outgrowth analysis. After one day of outgrowth at 37°C with 5% CO2, the cell count was measured using the PrestoBlue™ Cell Viability Reagent (Invitrogen™ #A13261) following the manufacturer’s protocol. The absorbance was then recorded at 560 nm excitation and 590 nm emission using a microplate reader (BioTeck).

### Western botting

Protein samples were subjected by SDS-PAGE and transferred onto nitrocellulose membranes for immunoblotting. Blots were blocked with 5% milk/BSA in TBS/0.1% Tween 20. For Snf1/AMPK phosphorylation analysis, blots were probed with anti-phospho-Thr172-AMPKα (1:1000 dilution; 40H9; Cell Signaling Technology). For total Snf1 and AMPK analysis, blots were probed with anti-MYC-Tag antibody (1:1000 dilution; #2272; Cell Signaling Technology) and anti-AMPKα antibody (1:1000 dilution; D5A2; Cell Signaling Technology), respectively. For Nrf2, p21 Waf1/Cip1 and β-Actin, blots were probed with anti-Nrf2 antibody (1:2000 dilution; E3J1V; Cell Signaling Technology), anti-p21 Waf1/Cip1 antibody (1:2000 dilution; E2R7A; Cell Signaling Technology), and anti-β-Actin antibody (1:2000 dilution; D6A8; Cell Signaling Technology), respectively. All blots followed by anti-rabbit IgG HRP secondary antibody (1:5000; NA934V, GE Healthcare). TORC1 activity experiments were performed as previously described ^71^. Phosphorylation of Sch9 was monitored by western blotting probed with anti-HA 3F10 antibody (1:2000; Roche Life Science, USA) followed by goat anti-rat HRP-conjugated antibody (1:5000; Santa Cruz Biotechnology). Blots were developed with ECL prime western blotting detection reagent (Amersham Pharmacia Biotech, USA) and quantified using ImageJ for the iBright CL1500 Imaging System (Thermo-Scientific).

### ATP analysis

Yeast cells were combined with a final concentration of 5% trichloroacetic acid (TCA) and subsequently placed on ice for a minimum of 5 minutes. Following this, the cells were washed, suspended in 10% TCA, and subjected to lysis using glass beads in a bead beater to facilitate ATP extraction. The ATP levels were quantified using the PhosphoWorks™ Luminometric ATP Assay Kit (AAT Bioquest) and then normalized based on the protein content, which was measured using the Bio-Rad protein assay kit.

### Quantitative real time PCR

Total RNA was extracted from yeast cultures treated under various conditions through mechanical lysis, following the previously described method ^71^, utilizing the manufacturer’s disruption protocol. For both yeast and human cells, total RNA extraction was conducted using the Qiagen RNeasy mini kit. RNA concentration and integrity were subsequently evaluated using the ND-1000 UV-visible light spectrophotometer from Nanodrop Technologies. Quantitative real-time PCR (qRT-PCR) experiments were carried out according to established protocols, employing the QuantiTect Reverse Transcription Kit (Qiagen) and the SYBR Fast Universal qPCR Kit (Kapa Biosystems). Gene abundance was determined in relation to the reference transcripts: *ACT1* for yeast and *GAPDH* for human cells.

### Quantification and statistical analysis

Statistical analysis of results such as mean value, standard deviations, significance, and graphing were performed using GraphPad Prism v.10 software. The comparison of obtained data was statistically performed using the Student’s t-tests, Ordinary One-way ANOVA and Two-way ANOVA followed by multiples comparison tests. In all the graph plots, P values are shown as *P < 0.05, **P < 0.01, ***P < 0.001, and ****P < 0.0001 were considered significant. ns: non-significant.

## AUTHOR CONTRIBUTIONS STATEMENT

YZ, AN, NAF, SY, TINC, JLJ, IWHJMG and CJ performed the experiments. MA designed, supervised the study and wrote the manuscript.

## ACKNOWLEDGMENTS

Ingrid Wen-Hui Jeanette Morel Gan and Chen Junqi would like to express their gratitude to the A*STAR Graduate Academy (AGA) for awarding them the A*STAR Research Internship Award (ARIA), which provided them with the valuable opportunity to work in Mohammad Alfatah’s Lab for Aging Biology Intervention Research (ABIR) at the Bioinformatics Institute (BII), ASTAR, Singapore. We thank Frank Eisenhaber for backing this research.

## SUPPLEMENTARY FIGURE LEGENDS

**Figure S1.**
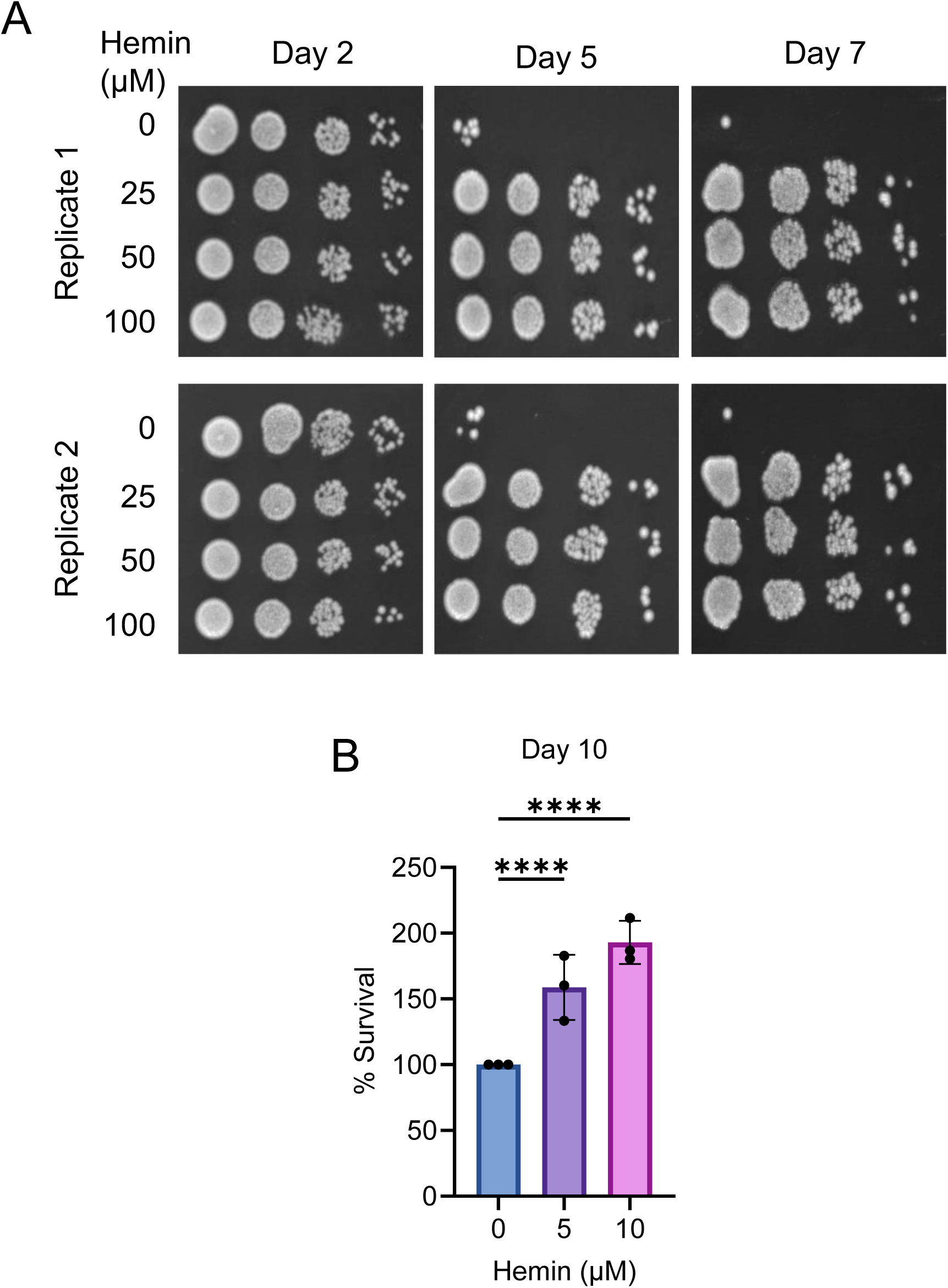
Chronological lifespan (CLS) analysis of wild-type yeast cells treated with hemin, Related to Figure 1. (A) Chronological lifespan (CLS) analysis was conducted for wild-type cells exposed to various concentrations of hemin in flasks. To initiate outgrowth, aging cells were serially diluted ten-fold and plated onto YPD agar. The data represents two experimental replicates. (B) The CLS of both wild-type cells with indicated concentrations of hemin was evaluated in SD medium using a 96-well plate format. The survival of aging cells was quantified through the PICLS method, employing a PI fluorescence-based assay. Measurements of PI fluorescence (excitation at 535nm, emission at 617nm) and OD600nm were obtained using a microplate reader. Fluorescence intensity values were normalized to OD600nm readings. The data are presented as means ± SD (n=3). Statistical significance was determined as follows: ****P < 0.0001 based on an ordinary one-way ANOVA followed by Dunnett’s multiple comparisons test.

**Figure S2.**
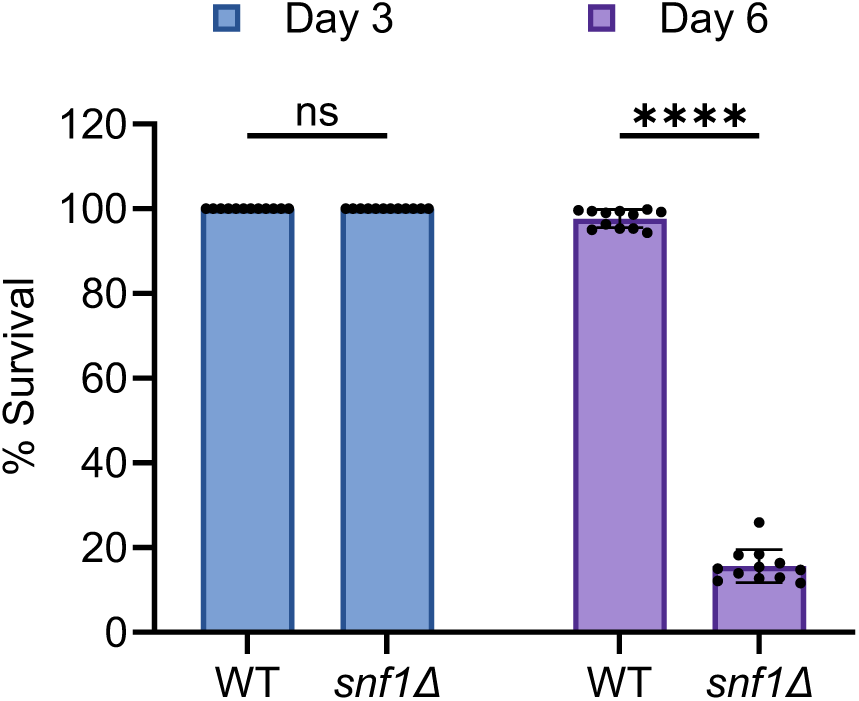
Chronological lifespan (CLS) analysis of wild-type and *snf1Δ* deletion yeast cells, Related to Figure 2. The chronological lifespan (CLS) of wild-type and *snf1Δ* deletion yeast cells was evaluated in SD medium using a 96-well plate. The survival of aging cells was measured at specific time intervals relative to the outgrowth observed on day 3. The data is presented as means ± SD (n=12). Statistical significance was determined as follows: ****P < 0.0001 and ns: non-significant, based on a two-way ANOVA followed by Šídák’s multiple comparisons test.

**Figure S3.**
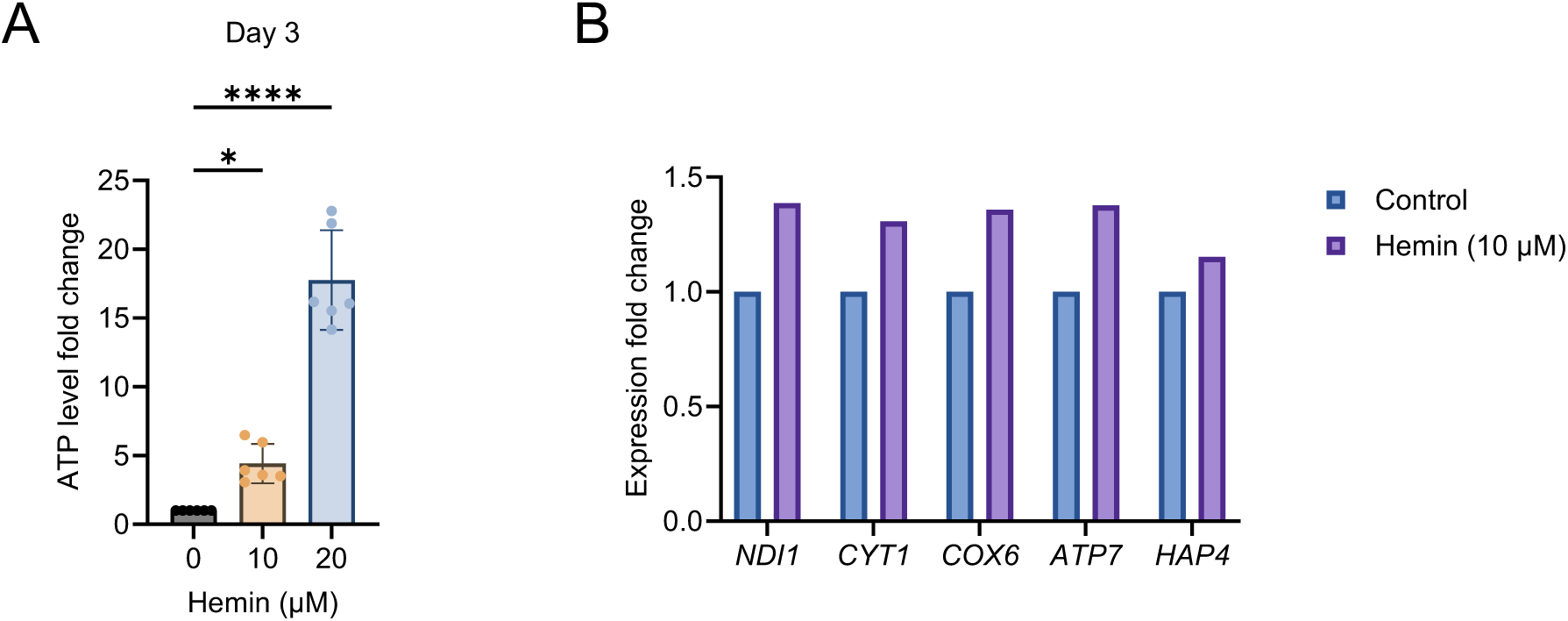
ATP and mitochondrial gene expression analysis of wild-type strain treated with hemin, Related to Figure 3. (A) ATP analysis of wild-type yeast cells incubated with different concentrations of hemin for 3 days. The data are presented as means ± SD (n=6). Statistical significance was determined as follows: *P < 0.05, and ****P < 0.0001, based on an ordinary one-way ANOVA followed by Dunnett’s multiple comparisons test. (B) Expression analysis of mitochondrial ETC genes by qRT-PCR in yeast cells incubated with 100 µM hemin for 6 hours, starting at ∼0.2 OD600 nm. The data are presented as means ± SD (n=2).

**Figure S4.**
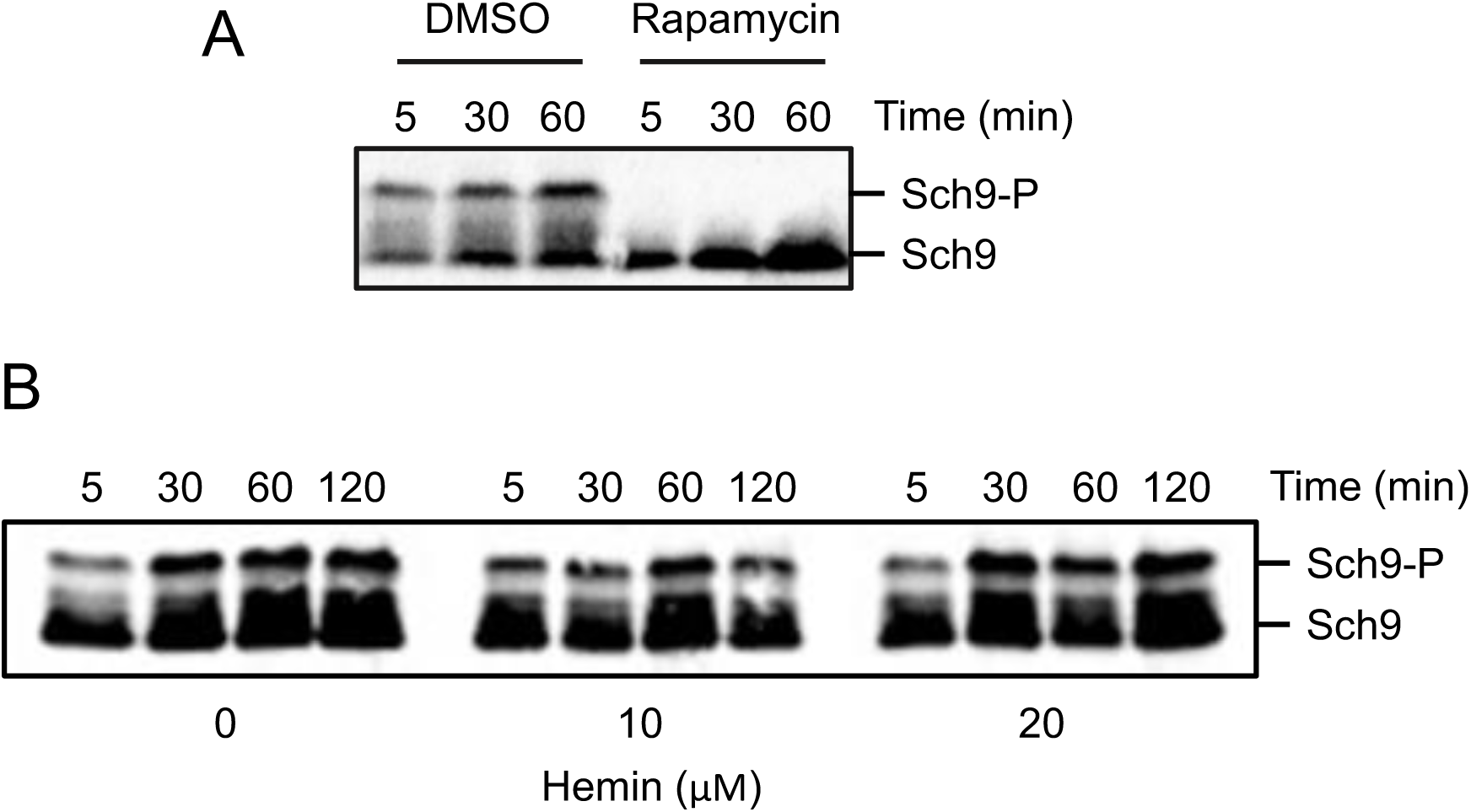
TORC1 activity analysis of wild-type strain (Sch9-6xHA-Tag) treated with rapamycin and hemin, assessed through Sch9 phosphorylation, Related to Figure 4. (A) Exponential cultures were treated with DMSO control and rapamycin (200 nM). Aliquots of the cultures were collected at indicated time points and utilized for preparing protein extracts. Sch9 phosphorylation was monitored through western blotting. (B) Exponential cultures were treated with varying concentrations of hemin. Aliquots of the cultures were collected at specified time intervals and used for preparing protein extracts. Sch9 phosphorylation was monitored via western blotting.

**Figure S5.**
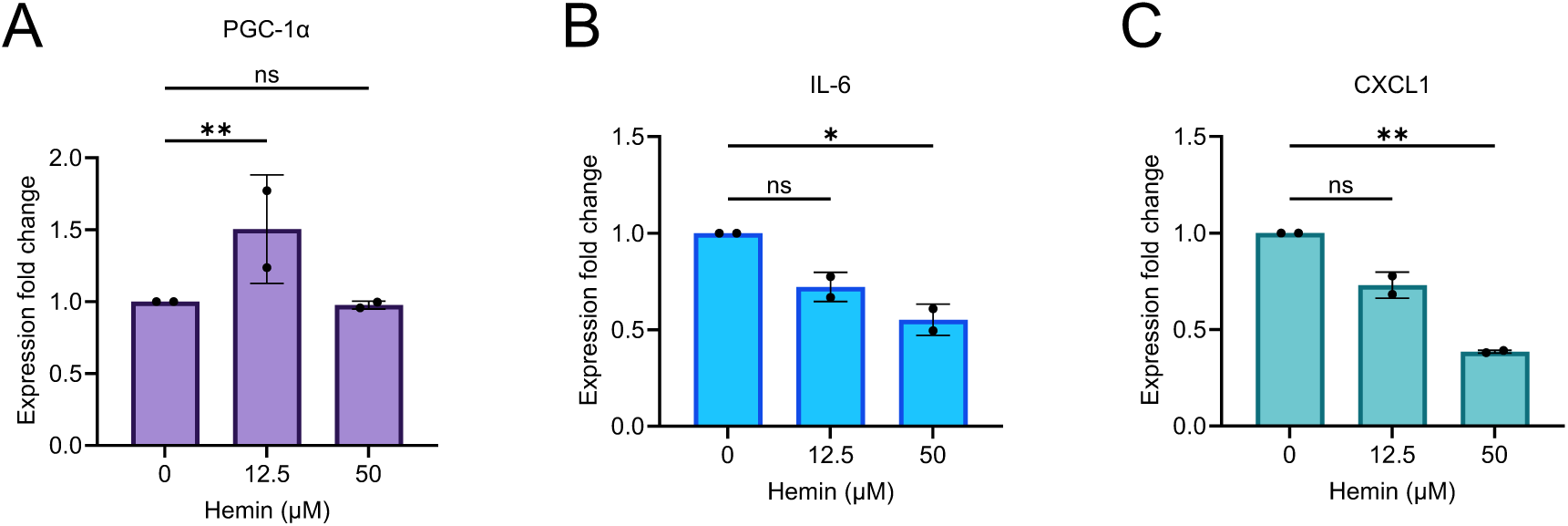
Gene expression analysis of human cells treated with hemin, Related to Figure 5. (A-C) Expression analysis of gene (A) PGC-1α, (B) IL-6 and (C) CXCL1 by qRT-PCR in human lung fibroblast cells (IMR90) treated with indicated concentrations hemin for 24 hours. The data are presented as means ± SD (n=2). Statistical significance was determined as follows: *P < 0.05, **P < 0.01, ns: non-significant based on a two-way ANOVA followed by Dunnett’s multiple comparisons test.

